# The Shh/Gli3 gene regulatory network precedes the origin of paired fins and reveals the deep homology between distal fins and digits

**DOI:** 10.1101/2020.09.08.287532

**Authors:** Joaquín Letelier, Silvia Naranjo, Ismael Sospedra, Javier Lopez-Rios, Juan Ramón Martinez-Morales, Neil Shubin, José Luis Gómez-Skarmeta

**Affiliations:** Centro Andaluz de Biología del Desarrollo (CABD), Consejo Superior de Investigaciones Científicas/Universidad Pablo de Olavide/Junta de Andalucía, Sevilla, Spain; Center for Integrative Biology, Facultad de Ciencias, Universidad Mayor, Santiago, Chile; Department of Organismal Biology and Anatomy, University of Chicago, Chicago, IL 60637

## Abstract

One of the central problems of vertebrate evolution is understanding the relationship among the distal portions of fins and limbs. Lacking comparable morphological markers of these regions in fish and tetrapods, these relationships have remained uncertain for the past century and a half. Here we show that *Gli3* functions in controlling the proliferative expansion of distal progenitors are shared among median and paired fins as well as tetrapod limbs. Mutant knockout *gli3* fins in medaka (*Oryzias latipes*) form multiple radials and rays, in a pattern reminiscent of the polydactyly observed in *Gli3* null mutant mice. In limbs, *Gli3* controls both anterior-posterior patterning and cell proliferation, two processes that can be genetically uncoupled. *In situ* hybridization, quantification of proliferation markers, and analysis of regulatory regions reveal that in paired and median fins, *gli3* plays a main role in controlling proliferation but not in patterning. Moreover, *gli3* downregulation in *shh* mutant fins rescues fin loss in a manner similar to how *Gli3*-deficiency restores digits in the limbs of *Shh* mutant mouse embryos. We hypothesize that the *Gli3/Shh* pathway preceded the origin of paired appendages and was originally involved in modulating cell proliferation. Accordingly, the distal regions of median fins, paired fins, and limbs retain a deep regulatory and functional homology that predates the origin of paired appendages.

---

A fundamental issue in vertebrate biology is the relationship between fins and limbs. Limbs have a characteristic skeletal pattern of stylopod, zeugopod and autopod, with a distal region composed of digits and mesopodial bones. While fossil taxa reveal intermediates in these conditions^1–3^, the appendages of extant actinopterygians lack common features that allow comparison among the distal regions. Teleosts, for example, have fins with both endochondral and dermal bones. The distal fin typically has a series of cartilage elements with no obvious homology to digits or mesopodials while the rays develop as dermal, not endochondral bones. Despite these dramatic differences in the distal anatomy of limbs and fins, the terminal region of fins reveals molecular similarities to limbs. Limbs have a characteristic pattern of two phases of expression of *Hox* genes, an early one that is associated with the specification of the stylopod and zeugopod and a later one associated with the formation of digits. Recent studies of the pattern of expression, function, and cell lineage of *Hox*-expressing cells in fish reveals that they have late phase activity that is comparable to that of limbs^4^. Lacking, however, is a knowledge of how deep these homologies extend in the tree of life. Are these kinds of similarities common to paired fins or are they a property of diverse paired and unpaired appendages of gnathostomes?

*Gli3*, a transcription factor expressed from early to late phases of limb development, has been shown to play multiple roles during limb morphogenesis. *Gli3* acts to set up the anterior-posterior axis of the limb bud and restricts *Shh* pathway activation to the posterior margin of the distal appendages^5–8^. This activity is mediated by the interactions of *Gli3* with *Hand2* and *Hoxd* transcription factors. *Gli3*-deficient mice show *Shh* pathway de-repression in the anterior limb bud margin, leading to the anterior expansion of the expression domains of posterior markers and concordant downregulation of anterior transcription factors^9–14^. Morphologically, these mutants reveal polydactylous manus and pes along with soft tissue fusion of digits. Most importantly, mouse limb buds lacking both *Shh* and *Gli3* have identical gene expression patterns to those of *Gli3* single mutants and their limb skeletons are indistinguishable^11,13^. More recent studies have revealed that the roles played by *Gli3* in patterning and cellular proliferation during appendage morphogenesis can be genetically uncoupled. The control of the proliferative expansion of the autopod progenitors is hence key to constrain the number of digits to the pentadactyl pattern^15^.

While *Gli3* functions and interactions with *Shh* are essential features of limb development, little is known about its role in fins. This deficit is unfortunate because an understanding of this issue could reveal the origin of distal patterns between the two organs. To address this, we generated *gli3* knockout mutants in medaka, a teleost fish with a single copy of *shh*, to assess gene expression, regulation and ultimately evolution of the morphoregulatory mechanisms mediated by *Gli3* in vertebrate appendages.

## RESULTS

To explore the role of *gli3* in medaka fin patterning, we deployed CRISPR-Cas9 to disrupt its coding region via an 86bp deletion in exon 5 (Fig. 1a, Extended Data Fig. 1). This mutation generates a frame shift that truncates the protein in the middle of the repressor domain. Adult fins could not be analyzed, as fish homozygous for this *gli3* inactivated allele die between 2 and 5 weeks of age, probably reflecting the pleiotropic functions of *gli3* in multiple tissues. By weeks 3-5, however, multiple anomalies were observed in the pectoral fin skeleton of *gli3* mutant escapers. In particular, these fins had an expanded number of radials and rays (Fig. 1b, c, e, f). Transient *gli3*-mutants raised to adulthood displayed greatly expanded fins, with multiple supernumerary elements (Fig. 1d, g, i, j). Hence, *gli3*-deficient fish are similar to mouse and human *Gli3* polydactylous mutants in having expanded appendages with extra bones.

**Figure 1.**
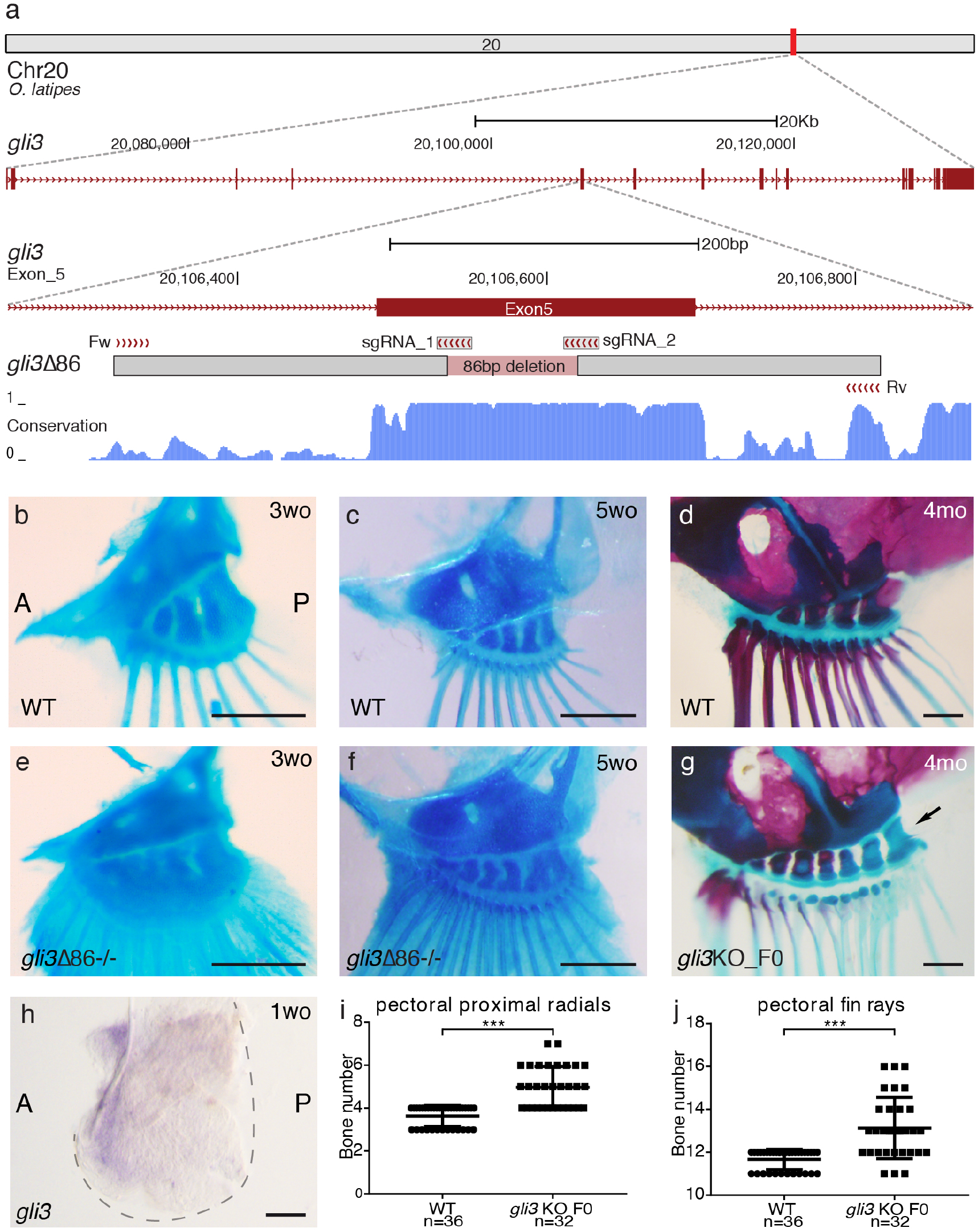
Medaka *gli3* mutants show an increased number of pectoral fin skeletal elements. a. Stable lines harboring Δ86 deletions in medaka *gli3* exon 5 were generated by CRISPR/Cas9. The diagram shows the position of the deletion relative to the single guide RNAs (sgRNAs) and primers used for screening along with the box of conservation with 4 fish species (stickleback, fugu, tetraodon and zebrafish). The Δ86 deletion results in the generation of a premature STOP codon that truncates the predicted protein upstream of the ZnFn DNA-binding domain of *Gli3*. b-g. Alcian blue and alizarin red staining of pectoral fin skeletons reveals a significantly increased number of proximal radial bones and fin rays in stable (e, f) and transient (g) *gli3* mutants. For b and c panels, n≥14 fins; for e and f panels, n≥12 fins. Scale bars 250*μ*m. h. Wild type pectoral fin showing the expression of *gli3* in the anterior region of the developing fin (n=10). Scale bar 100 *μ*m. i-j. Quantification of skeletal elements in adult (4m.o.) *gli3* crispants. Each point in the graphs represents the measurement of bone number in a single fin (n=36 WT, n=32 gli3 KO F0). An unpaired t-test was used for the statistical analysis of skeletal element number. \*\*\**P* = 5.06 × 10^-10^ for the comparison between WT (mean = 3.639) and *gli3* crispants (mean = 4.969) pectoral proximal radials bones. \*\*\**P* = 2.33 × 10^-7^ for the comparison between WT (mean = 11.67) and *gli3* crispants (mean = 13.13) pectoral fin rays. Bone staining procedures in juvenile (b, c, e, f) and adult fish (b, d) were performed in three independent experiments. A, anterior; P, posterior.

Next, we examined the expression profiles of genes involved in appendage patterning, in particular those known in tetrapods to interact with *Gli3* during limb bud outgrowth. Similar to tetrapod limbs, *gli3* expression in medaka is higher in the anterior margin of the pectoral fin bud (Fig. 1h). In mouse limb buds, *Gli3* inactivation causes the anterior expansion of posterior markers such as *Hand2*, 5’*Hoxd* and *Grem1*, and downregulation of anteriorly expressed genes (e.g., *Pax9*; Fig. 2a)^9,12–14^. In contrast, *in situ* hybridization analysis in medaka *gli3* mutant pectoral fins at stage 36 (dpf 6) revealed no obvious changes in the expression domains of these genes, as their expression patterns appear largely similar to those of wildtype embryos (Fig. 2b). Because *Gli3* has also been shown to control the proliferative expansion of mouse autopod progenitors through the direct transcriptional modulation of cell cycle regulators, we next used RT-qPCR to assess the expression levels of *Gli3-Shh* proliferative targets in fin buds (Fig. 2c). This analysis revealed the upregulation of *ccnd1* and *ccnd2* in *gli3*-deficient fins, revealing a role for *Gli3* in controlling distal proliferation during fin outgrowth.

**Figure 2.**
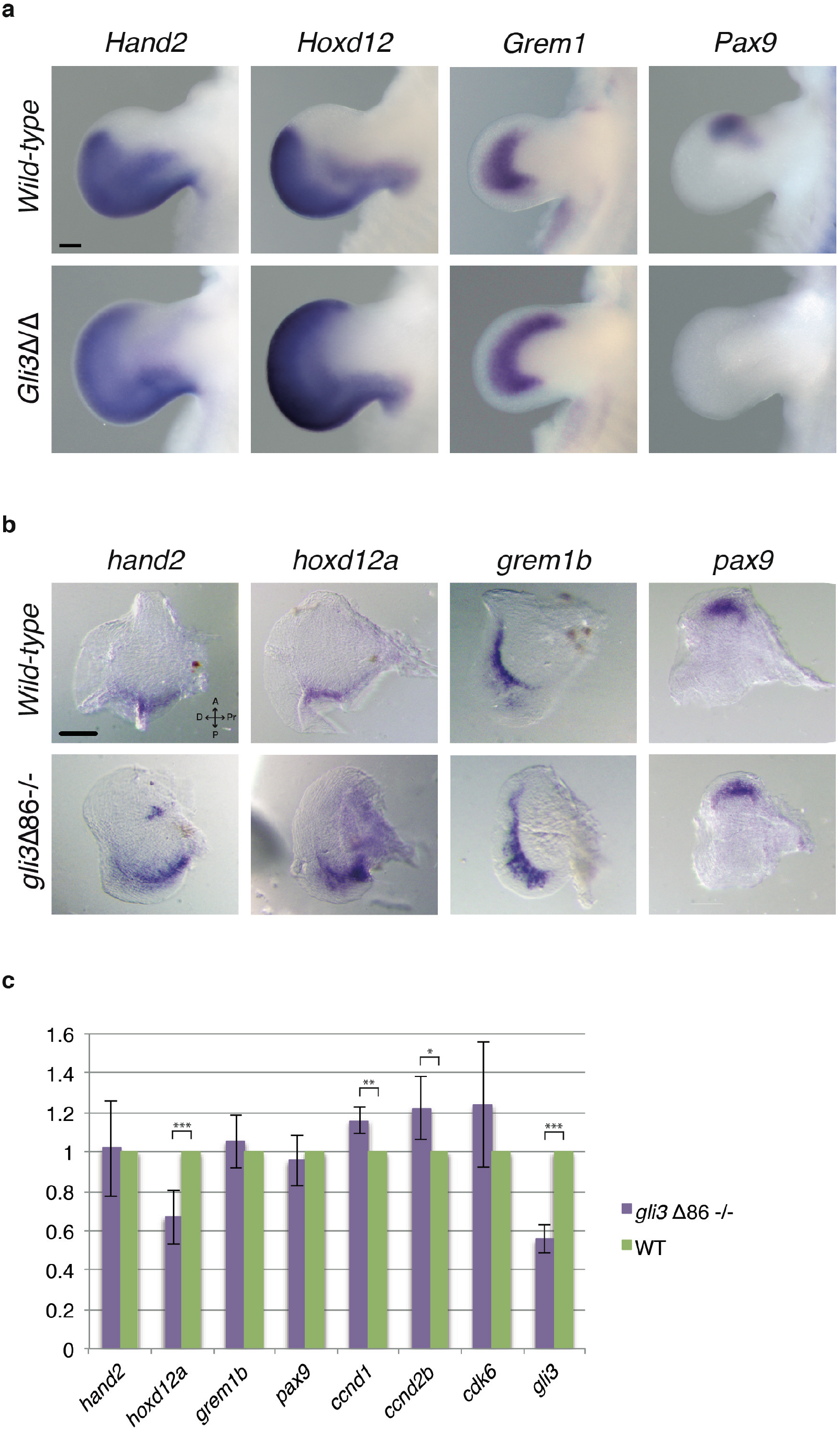
Expression of *gli3* target genes. a. *Hand2, Hoxd12* and *Grem1* expression is anteriorly expanded and *Pax9* expression lost in *Gli3*-deficient E11.5 mouse limb buds (n=3 per marker and genotype). Scale bar: 200 *μ*m. b. *In situ* hybridization assays in WT and *gli3*Δ86-/- medaka pectoral fins. At 6 dpf (stage 36), the inactivation of *gli3* does not greatly affect the expression pattern of *hand2, hoxd12a, grem1b* or *pax9*, all of them genes involved in limb patterning. A, anterior; P, posterior, Pr, proximal; D, distal. (n≥4 per marker and genotype). Scale bar: 100 *μ*m. b. Gene expression quantification by qPCR in WT and *gli3*Δ86-/- medaka pectoral fins at 11 dpf. The relative expression of the proliferation regulators *ccnd1* and *ccnd2b* is significantly increased in mutant fins. Mutant values (purple bars) are normalized against WT values (green bars), and represented as mean ± SD. Note that the *gli3*Δ86 allele is still transcribed in homozygous mutant fins, although at lower levels due to nonsense-mediated mRNA decay. \*\*\**P_gli3_* = 9.026 x 10^-9^, *P_hoxd12a_* = 0.0006; \*\**P_ccnd1_* = 0.0022; *P ≤ 0.05 *P_ccnd2b_* = 0.011.

Our gene expression and qPCR analyses point towards a role for *Gli3* in cell proliferation, rather than anterior-posterior patterning, during pectoral fin development. To explore this hypothesis, and reveal its phylogenetic distribution, we analyzed *Gli3* ChIP-seq data from mouse limb buds. This strategy revealed that *Gli3* preferentially binds at the promoters of these proliferation genes (Extended Data Fig. 2). These regulatory regions are present in diverse tetrapods and fishes, revealing their phylogenetic generality. In contrast, *Gli3* regulation of patterning genes (e.g., *Grem1* and *Hand2*), takes place mainly through distal *cis*-regulatory elements that are mostly not conserved in fish^16,17^. While this is clear for *Grem1*, in *Hand2* there are also *Gli3* binding sites at the promoter regions in mouse limb buds.

These data support the notion that the incorporation of *Gli3* in early patterning events in appendages is a more recent novelty within derived gnathostomes that evolved through the acquisition of new distal *cis*-regulatory elements (Extended Data Fig. 2).

A main function of *Shh* signaling is to antagonize the constitutive proteolytic processing of *Gli3* to its transcriptional repressor isoform^18^. Genetic analysis in mice has shown that *Gli3*-deficiency is able to rescue the loss of distal limb bones observed in *Shh* null embryos, leading to polydactyly that is identical to that observed upon *Gli3* inactivation alone^11,13^. To examine this relationship in medaka, we produced *gli3/shh* double mutants transiently, as *gli3*-deficient fish are not viable. Transient inactivation of *gli3* in a ZRS+sZRS *shh* mutant background^19^ is sufficient to rescue the *shh* loss of function phenotype (agenesis of pectoral, pelvic and dorsal fins;^19^). Analogous to the genetic interaction observed in mutant mouse autopods, the fin skeleton of ZRS+sZRS/*gli3* F0 double mutants resemble those of *gli3* crispants. This effect was seen in both paired and unpaired fins, as supernumerary bones were seen in pectoral, pelvic and dorsal fins (Fig. 3, Extended Data Fig. 3). Interestingly, as observed previously in ZRS+sZRS *shh* mutant^19^, the anal fin is not affected by any of these mutant conditions (Extended Data Fig. 4).

**Figure 3.**
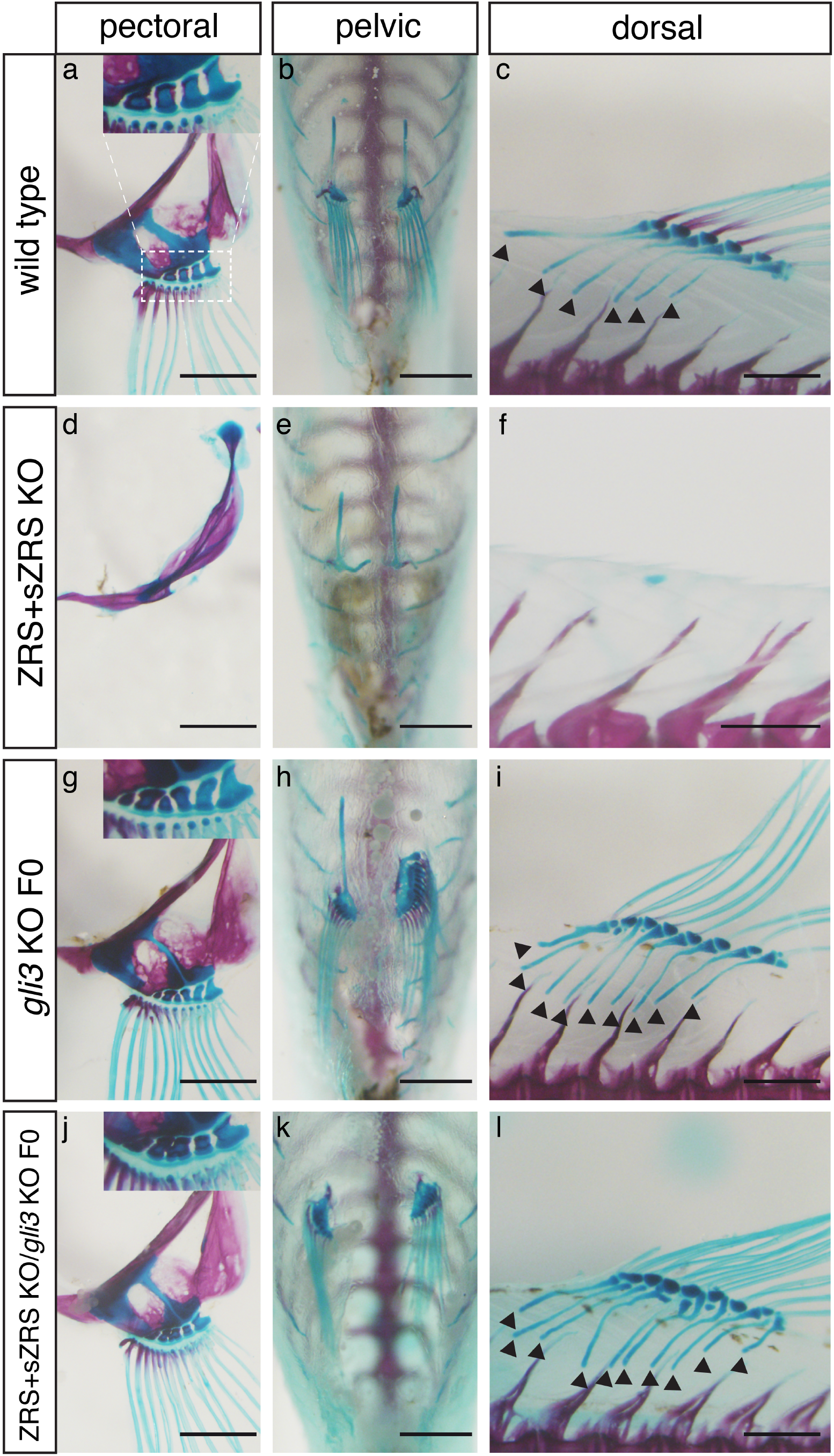
Mosaic inactivation of *gli3* in fins lacking *shh* completely rescues the formation of dorsal, pelvic and pectoral fins. a-l. Skeletal staining and fin morphology in adult fish. a-f. In contrast to WT (a-c), all fin elements are absent in the dorsal, pelvic and pectoral fins from ZRS+sZRS mutant fish (d-f). g-i. CRISPR/Cas9 disruption of *gli3* significantly increases the number of pectoral and dorsal fin bone elements (black arrowheads in i) compared to WT (a-c). j-l. *gli3* downregulation in the homozygous ZRS+sZRS *shh* mutant background totally rescues the dorsal, pelvic and pectoral fin phenotypes. Scale bars: 1mm. Microinjection of the *gli3* CRISPR cocktail and bone staining procedures were performed in three independent experiments.

## DISCUSSION

Altogether, our results show that the presence of the *shh/gli3* regulatory network in fish fins, so vital for limb formation and digit patterning, is primitive to limbs. Moreover, its functions in unpaired dorsal fins, widely recognized precursors of paired appendages, suggest that the recruitment of this network may have preceded the origin of paired fins themselves.

The correlation of expanded radials and rays in *gli3* fin mutants with the polydactyly in mouse *Gli3* mutants, points to a deep homology among the distal tissues of gnathostome appendages. Our analyses suggest that the primitive function of *Shh/Gli3* module in appendages was to control the proliferative expansion of the distal mesenchyme, while the anterior-posterior patterning systems controlled by *Gli3* likely evolved later during evolution, probably in the tetrapod lineage through the appearance of novel far-acting *cis*-regulatory regions^20,21^. Interestingly, the transcriptional control of cell cycle effectors is an ancient feature of HH signaling, as Ci, the fly ortholog of *Gli3*, also directly regulates several *cyclin* genes^22,23^.

Our results imply that the distal regions of appendages have a common evolutionary origin and that the *Shh/Gli3* network was modified in fish and tetrapod lineages to produce fin radials and rays in the former and digits in the latter. Interestingly, the only appendage which does not follow these rules, the anal fin, also has anomalous patterns of *Shh* regulation. The absence of these networks in anal fins points to a separate evolutionary origin for anal fins, presumably by independent cooption of fin patterning networks in a novel site.

## Extended data

**Extended Data Table 1 –** List of primers used in this study

**Extended Data Figure 1:**
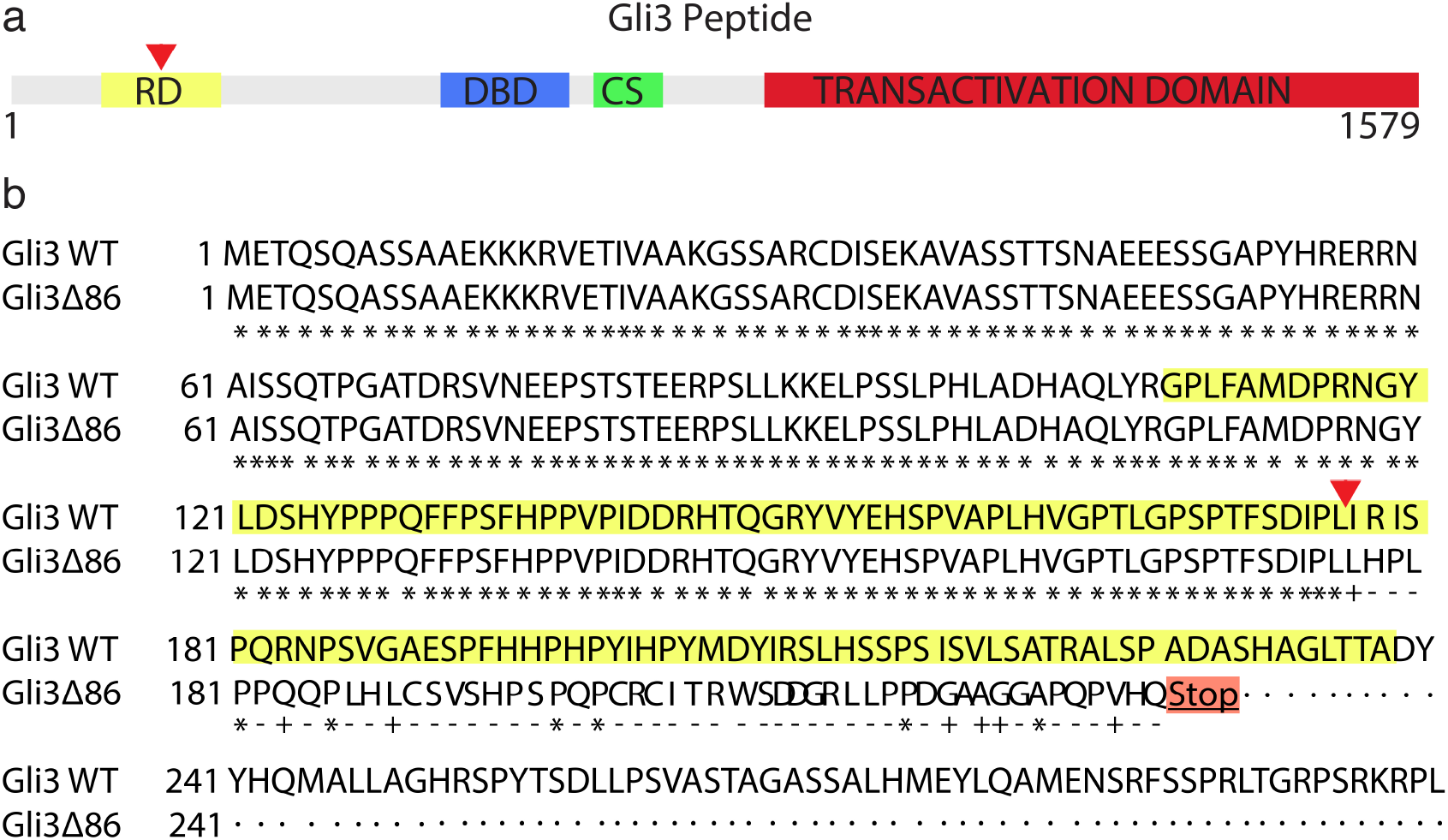
*gli3* Δ86 CRISPR/Cas9 induced mutation truncates the protein in the repressor domain. a. Scheme showing relevant domains of the Gli3 protein. RD: repressor domain (yellow), DBD: DNA binding domain (blue), CS: cleavage site (green) and transactivation domain (red). b. The Δ86 mutation (red arrowhead) generates a frame shift change in the middle of the repressor domain (yellow amino acid sequence) that results in a premature STOP codon that truncates the predicted protein upstream of the DBD.

**Extended Data Figure 2:**
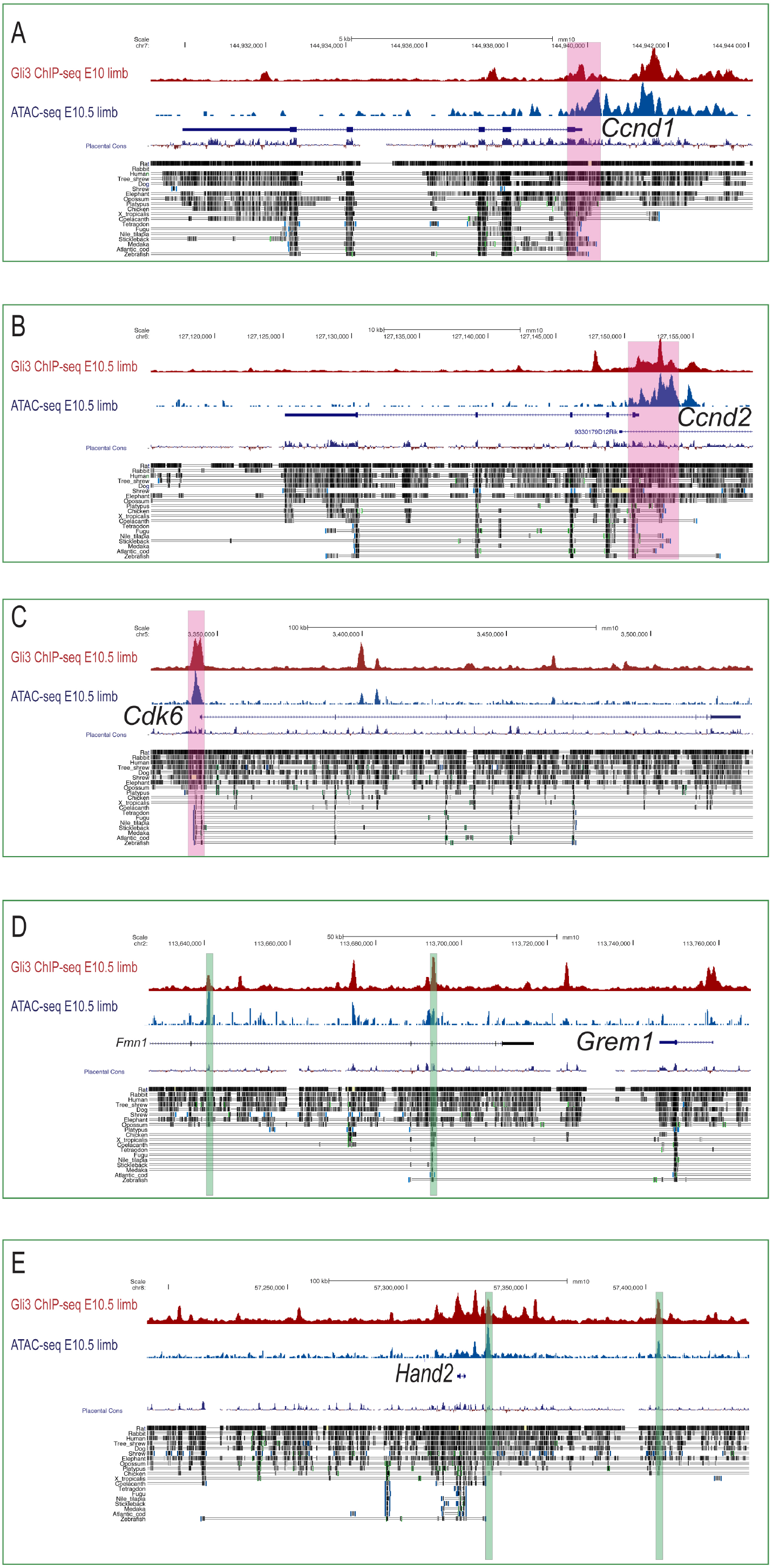
Regulatory landscapes of *Gli3* target genes. a-e. Red tracks show Gli3 ChIP-seq at E10.5 mouse limb buds^24^. Blue tracks show ATAC-seq data from E10.5 mouse limbs^25^. a-c. Cell cycle regulators (A, *Ccnd1;* B, *Ccnd2* and C, *Cdk6*), are mainly regulated by *Gli3* at the promoters through regions that are evolutionary conserved in fish genomes (highlighted in pink). d-e. Genes involved in limb patterning are mainly regulated by Gli3 through distal CREs, most of them not conserved in fish genomes. Green highlights show previously identified Gli3-bound limb regulatory elements^16,17^.

**Extended Data Figure 3.**
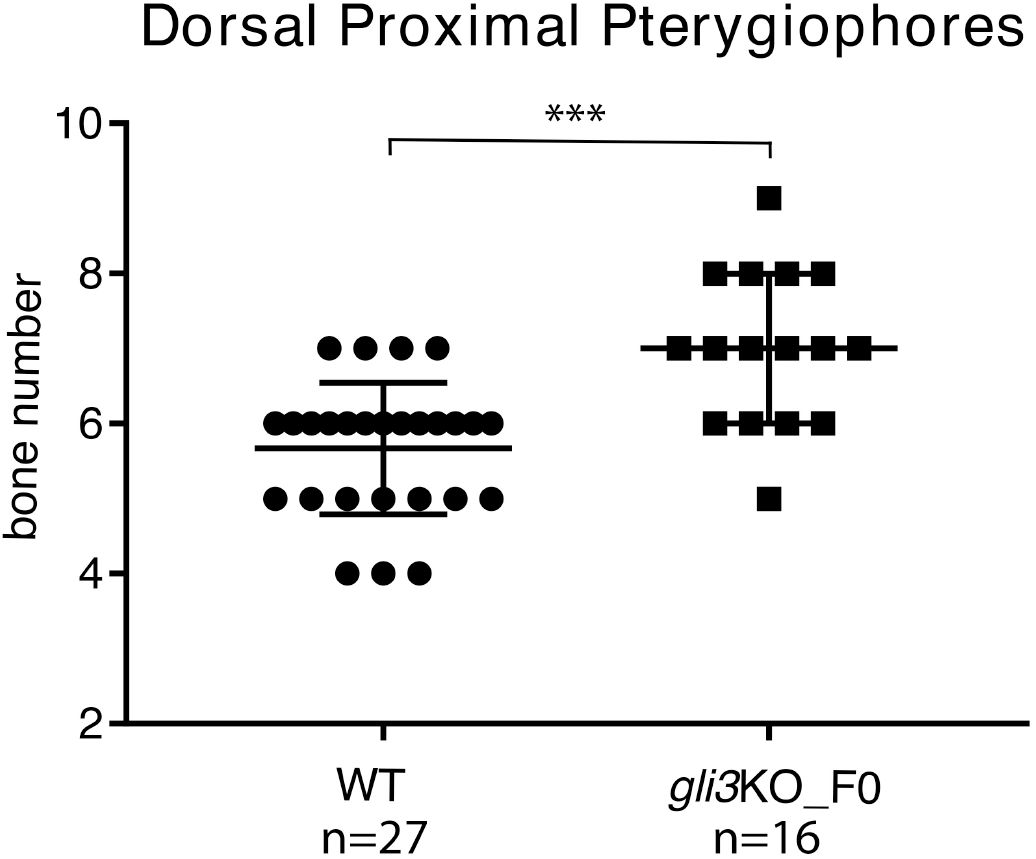
*gli3* crispants show increased bone number in the adult dorsal fin. a. Graph showing the number of proximal pterygiophore bones in the dorsal fin from WT and *gli3*_F0 crispants. Each point in the graph represents bone number in a single fin. An unpaired t-test was used for the analysis of bone quantification. \*\*\**P* = 5.33 × 10^-5^ for the comparison between WT (mean = 5.666) and *gli3* crispants (mean = 7). Microinjection of the *gli3* CRISPR mixture and skeletal staining were performed in three independent experiments.

**Extended Data Figure 4:**
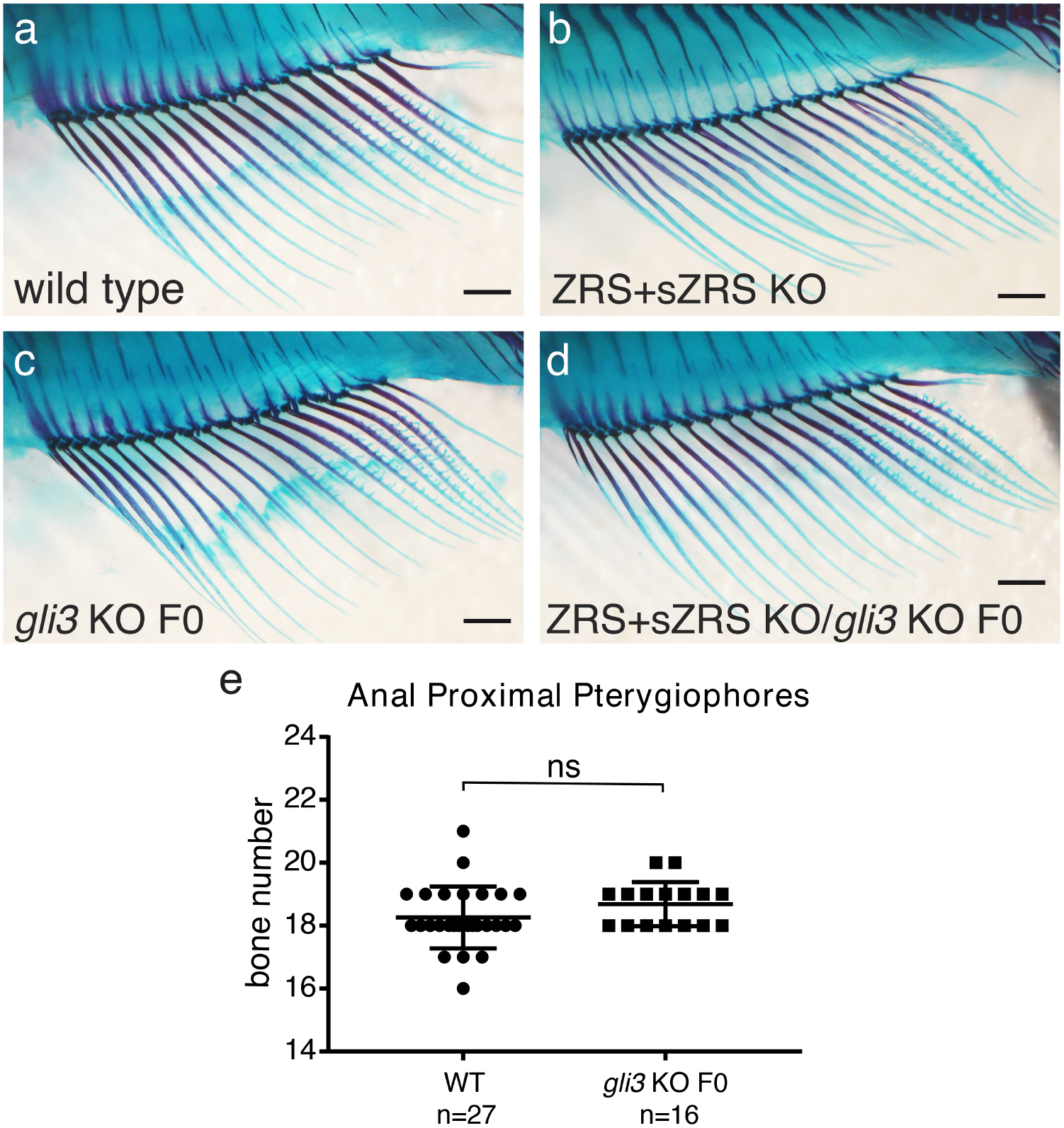
Anal fins are not affected in *gli3* and *shh* mutants. a-d. Skeletal staining showing adult (4m.o.) anal fins from WT (a), ZRS+sZRS KO (b), *gli3* F0 crispants (c) and compound *shh/gli3*_F0 mutants (d). Scale bars 1mm. e. No significant change in the number of anal proximal pterygiophores or fin rays (not shown) was observed between WT and *gli3* F0 crispants. Each point in the graph represents the measurement of bone number in a single fin (n=27 WT, n=16 *gli3* KO FO). An unpaired t-test was used for the statistical analysis of anal proximal pterygiophore bones. *P* = 0.1358 for the comparison between WT (mean = 18.26) and *gli3* crispants (mean = 18.69). All pictures are lateral views with anterior to the left. Microinjection of the *gli3* CRISPR cocktail and bone staining procedures were performed in three independent experiments. ns: not significant.

## Acknowledgements

We thank all members of JLGS laboratory for fruitful discussions. JLG-S has received funding from the European Research Council (ERC) under the European Union’s Horizon 2020 research and innovation programme (grant agreement No 740041), the Spanish Ministerio de Economía y Competitividad (grant BFU2016-74961-P) and was also supported, together with JL-R and JRMM by the institutional grant Unidad de Excelencia María de Maeztu (MDM-2016-0687 to the Department of Gene regulation and morphogenesis of Centro Andaluz de Biología del Desarrollo). JL-R and JRMM are supported by the Spanish Ministerio de Economía y Competitividad (grants BFU2017-82974-P and BFU2017-86339-P respectively). JL is financed by the Chilean FONDECYT agency (grant #11180727).

## Author contributions

JL, SN and IS designed and performed experiments. JL-R and JRMM conceived some experiments. NS and J.L.G.-S. coordinated the project, contributed to the study design and wrote the main text with input from all authors.

## Competing interests

The authors declare no conflict of interests.

Correspondence and requests for materials should be addressed to JL or JLG-S.

## METHODS

### Animal experimentation

All experiments involving fish and mice performed in this work conform European Community standards for the use of animals in experimentation and were approved by the Ethical Committees from the Universidad Pablo de Olavide, Universidad Mayor, Consejo Superior de Investigaciones Científicas (CSIC) and the Andalusian government.

### Fish stocks

Medaka wild-type (iCab) and ZRS+sZRS KO (Δ3.4kb^19^ strains were maintained and bred under standard conditions^26^. Embryos were staged in hours’ post-fertilization (hpf) as previously described^27^. Medaka *gli3*_Δ86 and ZRS_Δ3.4Kbs mutant alleles were maintained in heterozygosis due to the higher lethality of the homozygous mutants. In the case of *gli3*_Δ86-/- larvae, they die around three to four weeks post fertilization.

### Skeletal staining

Alcian blue and Alizarin red staining experiments were performed as previously described^19,28^. In brief, fish were fixed with 10% neutral-buffered formalin overnight or longer. After several washes with deionized water, cartilage was stained overnight using a 0.1% Alcian blue solution (Alcian Blue 8 GX, PanReac Applichem) in 30% acetic acid and 70% ethanol. Fish were then washed with deionized water and changed to a solution containing 1% trypsin from bovine pancreas (PanReac Applichem) and 30% saturated sodium borate for 8 hours (or longer) with gentle shaking at room temperature. After trypsin enzyme treatment, specimens were rinsed several times with deionized water and transferred to 0.5% aqueous KOH solution. Fish bones were finally stained overnight with a solution containing saturated Alizarin Red S (PanReac Applichem) in 0.5% KOH. After several washes with 0.5% KOH solution, fish were gradually transferred to glycerol for documentation. Specimens were visualized with and Olympus SZX16 binocular microscope and photographed with an Olympus DP71 camera.

### CRISPR/Cas9 design and *gli3* mutant generation

Two guided RNAs (sgRNAs) targeting exon5 of medaka *gli3* were designed using the CRISPRscan^29^ and CCtop^30^ CRISPR design online tools. sgRNAs were generated and purified for injection as previously described^31^. For sgRNAs generation the following primers were aligned (by PCR) to a universal CRISPR primer; *gli3* exon5 sgRNA1: 5’-taatacgactcactataGGGCGGATGTAGTCCATGTAgttttagagctagaa-3’ and *gli3* exon5 sgRNA2: 5’-taatacgactcactataGGGGTGAGATCCGAATGAGGgttttagagctagaa-3’ (in both primers the target site is identified by capital letters). Following synthesis, five nanoliters of a solution containing both sgRNAs at a concentration of 40 ng/*μ*L and Cas9 protein (Addgene #47327) at a concentration of 250 ng/*μ*L^32^ was injected into one-cell-stage medaka embryos. Oligos used for screening of genomic DNA deletions flanking CRISPR target sites were the following: forward primer 5’-CGTGAGTTTCACAGCAACAATTA-3’ and reverse primer 5’-CAGCCTCACTGATCAATTTCAG-3’. Mutations in *gli3* were analyzed by standard PCR and gel electrophoresis as the distance between both sgRNA PAM sequences was 82bp, long enough to create a deletion easily detected by a shift in the PCR band. Specific deletions in the *gli3* gene were further analyzed by Sanger sequencing of the PCR product from F1 embryos (Stabvida, Inc).

### Statistical analyses

The number of pectoral fin proximal radial bones, pectoral fin rays and dorsal proximal pterygiophores was manually counted in adult WT and *gli3* crispant fish after bone staining procedure. Differences in the number of skeletal elements between both groups were tested by applying an unpaired *t*-test using the GraphPad Prism software. For the statistical analysis of average differences in levels of expression of patterning and proliferation marker genes between mutant and WT samples with qPCRs, a paired two-tailed *t*-test was used.

### Medaka *in situ* hybridization

Depending on the genomic location of designed primers, antisense digoxigenin-labeled RNA probes were prepared from 4 dpf medaka cDNA or gDNA (see Extended Data Table1). *hand2, hoxd12a, grem1b* and *pax9*, RNA probes were synthetized by cloning the DNA amplified region in the Strataclone PCR Cloning Kit (240205-5, Agilent Technologies) (*hoxd12a*) or pGEM-T Easy Vector (A1360, Promega) (*pax9, grem1b, hand2*) and afterwards transcribing it to RNA from these vectors. The *gli3* probe was directly transcribed from the amplified DNA since the SP6-RNA Polymerase promoter sequence was included in the designed primers.

Heterozygous animals were mated in order to collect embryos to perform *in situ* hybridization (ISH) assays. The embryos were maintained at 28°C in E3 medium (NaCl 5mM, KCl 0,17 mM, CaCl_2_ 0,4 mM, MgMSO_4_ 0,16 mM and Methylene Blue 0,00003%) until 6 dpf, when they were dechorionated and fixated in PFA4% in PBS over 48 hours at 4°C. Finally, the embryos were dehydrated through an increasing MeOH washing series and kept at −20 °C until the experiments were performed. Overall, the specimens were prepared, hybridized, and stained as previously described in^33,34^. In particular, they were treated with 10 *μ*g of ProtK for 7 minutes for permeabilization and incubated with 2 ng/ *μ*l of each specific probe.

In order to analyze the expression of the different genes in mutant and WT siblings, samples were genotyped after performing *in situ* hybridization. For this, we extracted gDNA with Chelex 100 Sodium form (C7901, Sigma) from a piece of tail of single larvae and performed a standard PCR with primers flanking the deleted region (see Extended Data Table 1 and Fig. 1). Subsequently gene expression patterns in homozygous null and wild-type pectoral fins were analyzed and documented. Both fins of each genotyped larva were dissected and transferred to a drop of Methyl Cellulose 3% (M0387, Sigma-Aldrich) on a slide for documentation using an Olympus SZX16 (Model SZX2-ILLB) binocular scope and an Olympus DP71 camera.

### Quantitation of Transcript Levels in medaka fins by RT-qPCR

Larvae were raised as described above until 11 dpf and deeply anesthetized with 160 mg/l of Tricaine (Ethyl 3-aminobenzoate methanesulfonate salt, MS-222, Merck) before dissecting their pectoral fins. The dissection was performed in a drop of embryos’ medium and the fins were rapidly moved to an ice-cold drop of PBS. Every pair of dissected pectoral fins coming from a single larva was transferred to a separated 1.5 ml tube containing 50 *μ*l of TRIsure (BIO-38032, Bioline) and kept at −20°C in until larvae were genotyped (see *in situ* hybridization section). When 20 pairs of homozygous mutant and WT pectoral fins were collected, all the fins with the same genotype were pulled together and RNA was extracted following TRIsure manufacturer’s instructions.

The same amount of mutant and WT RNA was used to synthetize cDNA with High Capacity cDNA Reverse Transcript kit (Applied biosystems-Thermo Fisher Scientific, cat. 4368814). A total of four different samples (4 mutant and 4 WT) were prepared and a minimum of two were used to test the expression levels of the different targets.

The expression levels of medaka *hand2, hoxd12a, grem1b, pax9, ccnd1, ccnd2b, cdk6* and *gli3* in the developing fins were quantified through RT-qPCR (CFX96 real-time C1000 thermal cycler, Bio-Rad) and normalized to the expression level of the housekeeping gene *ef1a* (Extended Data Table1). qPCR reactions were performed in triplicates and repeated in a minimum of six different experiments from at least two different biological samples (described above) using iTaq Universal SYBR green Supermix (Bio-Rad, cat. 172-5124) according to the manufacturer’s instructions. The expression levels in mutant samples were calculated in relation to wild-type controls (average set to 100%). Assuming a normal distribution of the data, a paired two-tailed *t*-test was performed to test significance of differences among samples averages.

### Mouse experiments

E11.5 (44-48 somites) *Gli3*-deficient and wild-type embryos were processed for in situ hybridization with riboprobes recognizing *Hand2, Hoxd12, Grem1* and *Pax9* transcripts (n=3 per marker and genotype), as previously described^15^.

